# Fusogenic Structural Changes in Arenavirus Glycoproteins are Associated with Viroporin Activity

**DOI:** 10.1101/2023.02.20.529181

**Authors:** You Zhang, Joanne York, Melinda A. Brindley, Jack H. Nunberg, Gregory B. Melikyan

**Author notes:** Corresponding author (GBM).

## Abstract

Many enveloped viruses enter host cells by fusing with acidic endosomes. The fusion activity of multiple viral envelope glycoproteins does not generally affect viral membrane permeability. However, fusion induced by the Lassa virus (LASV) glycoprotein complex (GPc) is always preceded by an increase in viral membrane permeability and the ensuing acidification of the virion interior. Here, systematic investigation of “leaky” LASV fusion using single pseudoparticle tracking in live cells reveals that the change in membrane barrier function is associated with the fusogenic conformational reorganization of GPc. We show that a small-molecule fusion inhibitor or mutations that impair viral fusion by interfering with GPc refolding into the post-fusion structure prevent the increase in membrane permeability. We find that the increase in virion membrane permeability occurs early during endosomal maturation and is facilitated by virus-cell contact. This increase is observed using diverse arenavirus glycoproteins, whether presented on lentivirus-based pseudoviruses or arenavirus-like particles, and in multiple different cell types. Collectively, these results suggest that conformational changes in GPc triggered by low pH and cell factor binding are responsible for virion membrane permeabilization and acidification of the virion core prior to fusion. We propose that this viroporin-like activity may augment viral fusion and/or post-fusion steps of infection, including ribonucleoprotein release into the cytoplasm.

**Authors summary:** Fusion of enveloped virus with host cell membranes is mediated by extensive conformational changes in viral glycoproteins, triggered by binding to cognate receptors and/or by exposure to acidic pH in the maturing endosome. We have previously reported that, unlike many other viral glycoproteins, pH-triggered endosomal fusion by the Lassa virus viral glycoprotein complex (GPc) is preceded by a mild permeabilization of the viral membrane. Here, we provide evidence that this activity is associated with early fusogenic changes in the GPc and is markedly enhanced by virus-cell contact. Permeabilization is induced by the glycoproteins of diverse arenaviruses and occurs in multiple target cell types. Based on these observations, we propose that, by analogy to the influenza virus M2 channel, membrane permeabilization and the resultant acidification of the arenavirus interior facilitate viral fusion and/or post-fusion steps of infection.

## Introduction

Arenaviruses initiate infection by entering cells *via* endocytosis and fusion of their envelope membrane with the endosomal membrane. Arenavirus-cell fusion is mediated by the trimeric viral glycoprotein complex (GPc), each protomer of which consists of three noncovalently associated subunits, GP1, GP2, and a stable signal peptide (SSP) [1–5]. The membrane-distal GP1 subunit engages host attachment factors and receptors [4, 6–14], while conformational changes in the transmembrane GP2 subunit [15] drive the merger of viral and cellular membranes. These fusogenic changes in GPc are triggered by low endosomal pH and may be augmented by interactions with endosome-resident viral co-receptors. For example, the Old-World Lassa arenavirus (LASV) engages α-dystroglycan on the cell surface and, upon entry into acidic endolysosomes, switches to the LAMP1 receptor [16–19].

We have developed labeling and imaging strategies for robust single virion tracking and detection of individual HIV-1 pseudovirus fusion events. These strategies include incorporation of an HIV-1 protease-cleavable Vpr construct that produces free mCherry, a viral content marker, and a pH-sensitive YFP-Vpr protein packaged into the mature viral core, to detect changes in pH [20]. With this labeling approach, virus-cell fusion is manifested by a rapid release of mCherry through the fusion pore, whereas the viral core-associated YFP signal persists, aiding reliable detection of single virus fusion events [20]. Using this HIV-1 pseudovirus platform, we have studied entry driven by a variety of viral glycoproteins, including the Influenza A virus (IAV) HA, Vesicular Stomatitis Virus (VSV) G and Avian Sarcoma and Leukosis Virus (ASLV) Env [21–31], all of which mediate virus entry from acidic endosomes.

Recent investigations of the LASV entry mechanisms using HIV-1 pseudoparticles (LASVpp) revealed an unusual fusion phenotype. In contrast to “tight” fusion induced by other viral glycoproteins, LASVpp fusion is “leaky”, involving permeabilization of the viral membrane prior to fusion pore formation [31, 32]. The increase in membrane permeability to protons is detected as quenching of pH-sensitive YFP-Vpr fluorescence. Subsequent virus-endosome fusion results in a loss of the free mCherry signal due to its release into the cytoplasm, and simultaneous recovery of the YFP signal reflecting re-neutralization of the viral interior [31, 32]. Analysis of the temporal relationship between changes in the intraviral pH and mCherry release allowed us to delineate the initial dilation of fusion pores. We have demonstrated that binding to human LAMP1, while not strictly required for LASV fusion, strongly promotes the enlargement of the nascent fusion pore [32]. However, the mechanism of viral membrane permeabilization and the potential role of the virion interior acidification in LASV entry/infection remain unclear.

Here, using an HIV-1 based pseudovirus platform and arenavirus-like particles (VLPs), we demonstrate that viral membrane permeabilization is a common feature of fusion mediated by GPc of diverse arenaviruses and in different cell types. We also show that the increase in viral membrane permeability is mediated by GPc upon endosome acidification and is augmented by virus contact with the cell. Importantly, genetic, and pharmaceutical interventions that impair LASV GPc’s fusion activity diminish its ability to increase membrane permeability, suggesting that functional refolding of the arenavirus glycoprotein compromises the barrier function of the viral membrane. Together, our results support the notion that a “leaky” virion membrane and the resulting acidification of the virion interior prior to virus-cell membrane fusion may be important for post-fusion entry events.

## Results

### Viral membrane permeabilization is an intrinsic feature of arenavirus GPc-mediated fusion

We have previously reported that membrane fusion by Lassa GPc-pseudotyped HIV-1 particles (LASVpp) is associated with an increase in virion membrane permeability and acidification of the virion interior, regardless of whether membrane fusion is triggered by endosomal acidification or exposure to low pH at the cell surface [31, 32]. Permeabilization of the virion membrane consistently occurs prior to formation of the fusion pore. In these studies, membrane permeabilization and viral fusion events were detected by labeling the pseudovirus with mCherry-CL-YFP-Vpr, which is cleaved by the HIV-1 protease at the cleavage site (CL) to generate free mCherry, and with YFP-Vpr, which remains associated with the virion core (Fig. 1A) [20]. Permeabilization of the viral membrane and acidifications of the virion interior are manifested by quenching of the pH-sensitive YFP signal; free mCherry is too large to diffuse out of the permeabilized virions. Subsequent formation of the fusion pore leads to a quick recovery of YFP fluorescence due to re-neutralization of viral interior and loss of mCherry signal due to release into the cytoplasm (Fig. 1A, B and Movie S1) [20, 32]. This “leaky” phenotype contrasts with “tight” virus-endosome fusion by other viral glycoproteins that is not associated with YFP quenching (Suppl. Fig. S1 and Movie S2). It is unclear whether acidification of the virion interior occurs through defects forming in the viral lipid membrane or through a channel-like proteinaceous pore. We will therefore operationally refer to this phenomenon as viral membrane permeabilization.

**Figure 1.**
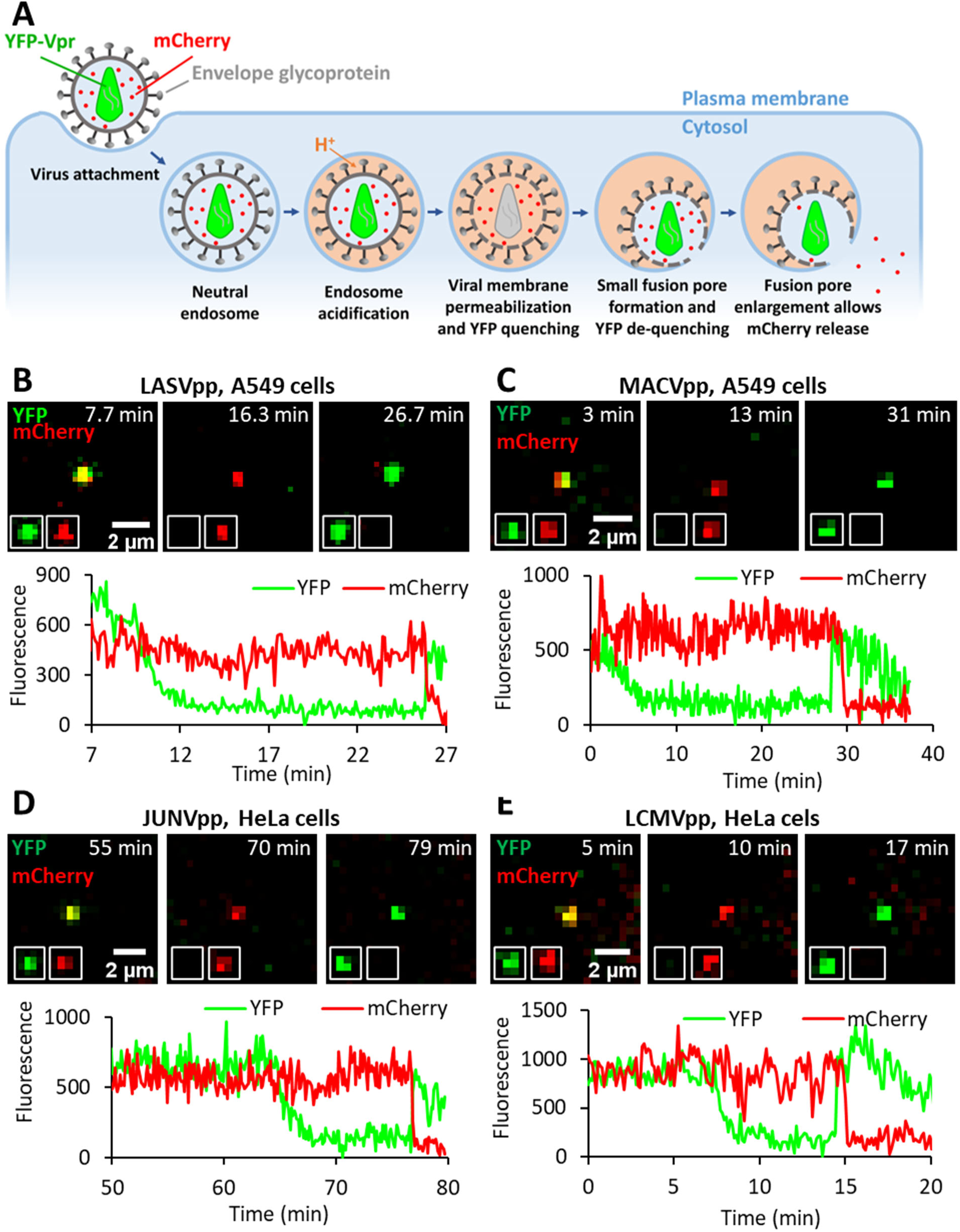
Arenavirus membrane permeability increases prior to virus fusion. (A) Illustration of mCherry-CL-YFP-Vpr labeled single LASVpp fusion. LASVpp is internalized and trafficked to acidic endosomes where the viral membrane is permeabilized. Increases in viral membrane permeability lead to acidification of the virus’ interior which is manifested in YFP signal quenching. LASVpp-endosome fusion results in mCherry release into the cytoplasm and concomitant re-neutralization of the virus’ interior, seen as recovery of YFP signal. (B) Single LASVpp fusion with A549 cell. Time-lapse images (top) and fluorescence traces (bottom) show virus interior acidification (YFP quenching) at 12.3 min and fusion (YFP dequenching and mCherry loss) at 25.9 min (see Movie S1). (C) A single MACVpp fusion event in A549 cell. Time-lapse images (top) and fluorescence traces (bottom) show virus interior acidification (YFP quenching) at 7.0 min and fusion (YFP dequenching and mCherry loss) at 29.0 min. (D) A single JUNVpp fusion event in HeLa cell. Time-lapse images (top) and fluorescence traces (bottom) show YFP quenching at 68.0 min and fusion (YFP dequenching and mCherry loss) at 77.1 min. (E) A single LCMVpp fusion event in HeLa cell. Time-lapse images (top) and fluorescence traces (bottom) show YFP quenching at 8.8 min and fusion (YFP dequenching and mCherry loss) at 14.9 min. A slightly delayed mCherry release after YFP dequenching in A and C is due to a slower dilation of fusion pores to sizes that allow mCherry release.

In order to determine the generality of GPc-mediated viral membrane permeabilization, we examined LASVpp entry into different cell types. Our results (Fig. S2) show that “leaky” LASVpp fusion is cell type-independent, as it consistently occurs in unrelated human and simian cells lines. The efficiency of “leaky” LASVpp fusion (percent of cell-bound particles that release mCherry after YFP quenching) varied depending on the cell line, with U2OS cells being the most permissive and VeroE6 cells being the least permissive to fusion (Fig. S2). Furthermore, single HIV-1 particles pseudotyped with the glycoproteins of another Old World arenavirus (LCMVpp) or distantly related New World arenaviruses (MACVpp and JUNVpp) also reproducibly undergo “leaky” fusion in acidic endosomes (Fig. 1C-E and Suppl. Fig. S2).

To exclude the possibility that “leaky” fusion might be an artefact of HIV-1 pseudotyping with arenavirus GPc, we generated arenavirus-like particles (VLPs) consisting of Candid-1 (JUNV vaccine strain [33, 34]) matrix (Z) protein and nucleoprotein (NP), bearing GPcs of diverse arenaviruses, essentially as described in [35]. In order to track VLPs and detect viral membrane permeabilization and fusion, we fluorescently labeled NP by replacing the D93 residue in an exposed loop [36] with the YFP sequence, for sensing intraviral pH, or with the pH-insensitive mCherry sequence (referred to as NP-DYFP and NP-DmCherry, respectively, Fig. 2A). Virions produced using a mixture of NP-DYFP and NP-DmCherry are positive for both markers. Unlike the HIV-1 based pseudoviruses labeled with cleavable mCherry-CL-YFP-Vpr (Fig. 1), the tagged NP proteins are not subject to cleavage and remain associated with the viral ribonucleoprotein complex (Fig. 2A). Using this VLP system, acidification of the virion interior and formation of the fusion pore are detected through YFP quenching and the subsequent recovery of the YFP signal upon re-neutralization (Fig. 2B-E). The pH-insensitive mCherry signal serves as reference for ensuring that YFP fluorescence changes are not due to particle departure from a focal plane.

**Figure 2.**
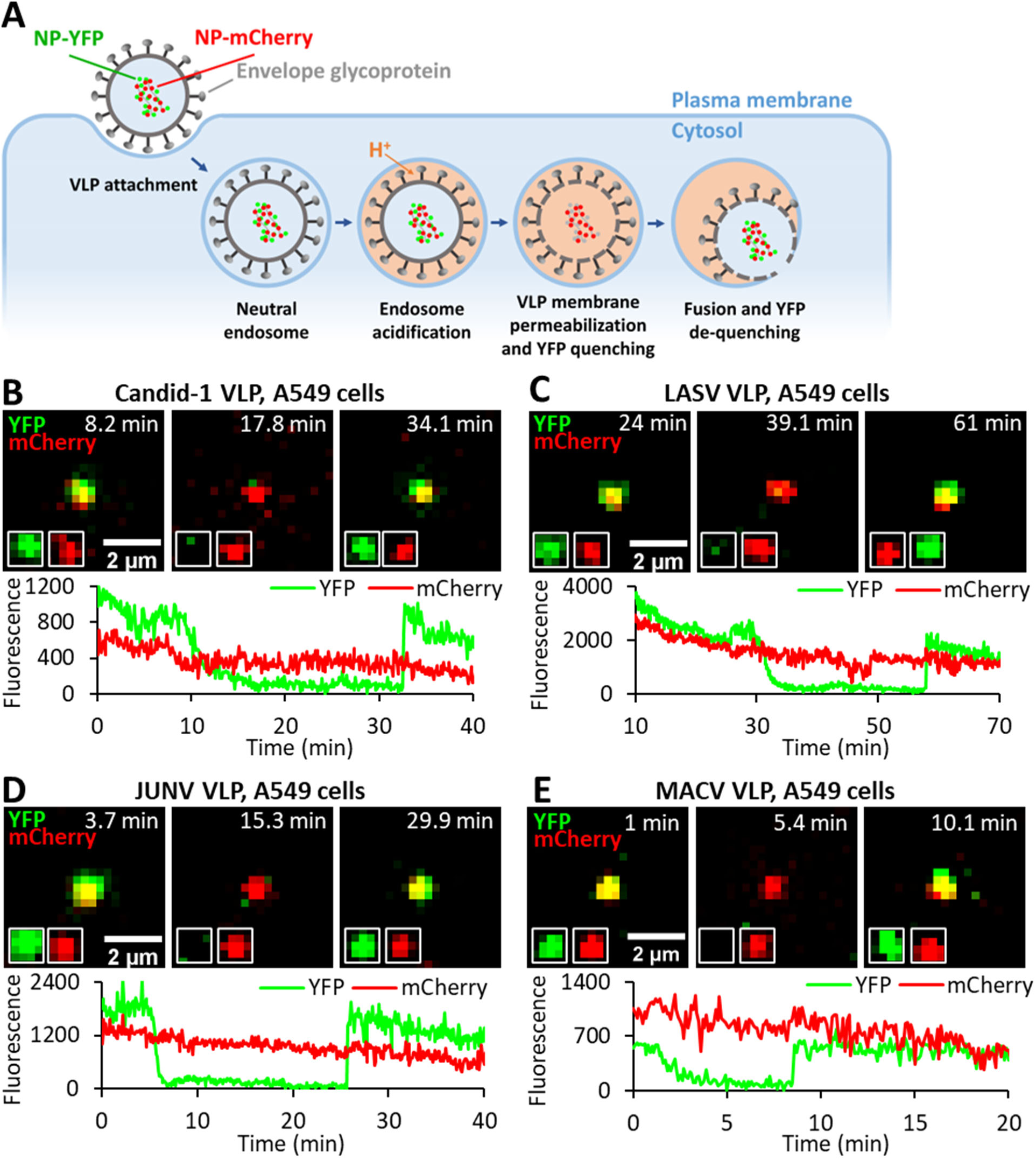
Arenavirus VLPs undergo membrane permeabilization prior to fusion. (A) Illustration of single VLP fusion labeled with NP-DmCherry/NP-DYFP. VLP is internalized, trafficked to acidic endosomes where the viral membrane is permeabilized, leading to acidification of the viral interior and quenching of YFP fluorescence. Subsequent fusion of VLP and endosomal membrane neutralizes the VLP interior and results in recovery of YFP signal. (B) A single Candid-1 VLP fusion event in A549 cell. Time-lapse images (top) and fluorescence traces (bottom) show VLP interior acidification (YFP quenching) at 12.1 min and fusion (YFP dequenching) at 33.4 min. (see Movie S3). (C) A single LASV VLP fusion event in A549 cell. Time-lapse images (top) and fluorescence traces (bottom) show virus interior acidification (YFP quenching) at 34.0 min and fusion (YFP dequenching) at 57.9 min. (D) A single JUNV VLP fusion event in A549 cell. Time-lapse images (top) and fluorescence traces (bottom) show VLP interior acidification (YFP quenching) at 7.3 min and fusion (YFP dequenching) at 25.6 min. (E) A single MACV VLP fusion event in A549 cell. Time-lapse images (top) and fluorescence traces (bottom) show virus interior acidification (YFP quenching) at 4.0 min and fusion (YFP dequenching) at 8.7 min.

This VLP labeling approach allows the detection of single fusion events mediated by Candid-1, JUNV, LASV and MACV GPcs (Fig. 2B-E). All arenavirus glycoproteins tested reproducibly increase the VLP membrane permeability, seen as NP-DYFP quenching, and mediate virus fusion with endosomes, as evidenced by NP-DYFP signal dequenching. Thus, viral membrane permeabilization prior to arenavirus GPc-mediated fusion pore formation is independent of the pseudotype platform, occurring in both HIV- and arenavirus-based particles. Quantitative differences in the efficiency and kinetics of VLP-cell fusion mediated by different GPcs (Suppl. Fig. S3) are likely caused by varied levels of expression of these glycoproteins or incorporation into Candid-1-based VLPs. The widespread occurrence of “leaky” fusion among arenaviruses suggests that membrane permeabilization and the resulting acidification of the virion interior may be an obligatory intermediate step of arenavirus GPc-mediated fusion.

### Permeabilization of arenavirus membrane occurs shortly after exposure to low pH

GPc-mediated membrane fusion can be triggered by exposure to acidic pH, independent of a cell surface receptor [7, 37]. To investigate whether cell contact is necessary for virion membrane permeabilization, we exposed coverslip-adhered LASVpp to a pH 5.0 membrane-impermeable citrate buffer. Only 3.9% of particles exhibited immediate YFP quenching upon acidification, demonstrating that nearly all pseudoviruses have intact membranes that limit the diffusion of protons (Suppl Fig. S4). To assess the temporal relationship between endosome acidification and LASVpp membrane permeabilization, pseudoviruses were co-labeled with two pH-sensitive fluorescent proteins – HIV-1 Gag-ecliptic pHluorin (Gag-EcpH, intraviral sensor) and pDisplay-pHuji, an external pH-sensor that is anchored to the virion membrane by the transmembrane domain of platelet-derived growth factor receptor (Fig. 3A) [38]. Pseudovirus entry into cells was synchronized by pre-binding dual-labeled particles to cells in the cold and quickly shifting to 37 °C. Upon endocytic uptake of the virus by the cell, endosomal acidification is manifested by a loss in the pHuji signal, whereas LASVpp membrane permeabilization is manifested by the loss of the EcpH signal (Fig. 3B). In contrast, HIV-1 particles pseudotyped with the Influenza virus HA (IAVpp) exhibited quenching of the surface marker (pHuji) but not the internal sensor (EcpH, Fig. 3C). Interestingly, despite the presence of GPc on virtually all pseudovirion particles (Suppl. Fig. S5), only 14% of cell-bound particles exhibited a loss of pHuji signal within 2 hrs of infection (Fig. 3D), suggesting that LASVpp entry into acidic compartments in A549 cells is inefficient and/or slow. As expected, loss of pHuji signal was potently inhibited by Bafilomycin A1 pretreatment, which blocks endosome acidification (Fig. 3D). Importantly, the kinetics of single particle pDisplay-pHuji and Gag-EcpH quenching were similar (half-times 55.2 and 59.8 min, respectively, Fig. 3E), implying that acidification of virus interior after exposure to low pH is faster than the time required for virus uptake and acidification of virus-carrying endosomes. Indeed, when the lag time between pHuji quenching (acidification of endosomal lumen) and subsequent loss of EcpH signal (acidification of the virion interior) was measured for individual viral particles, the average delay between endosomal acidification and viral membrane permeabilization was 0.8 min (Fig. 3F).

**Figure 3.**
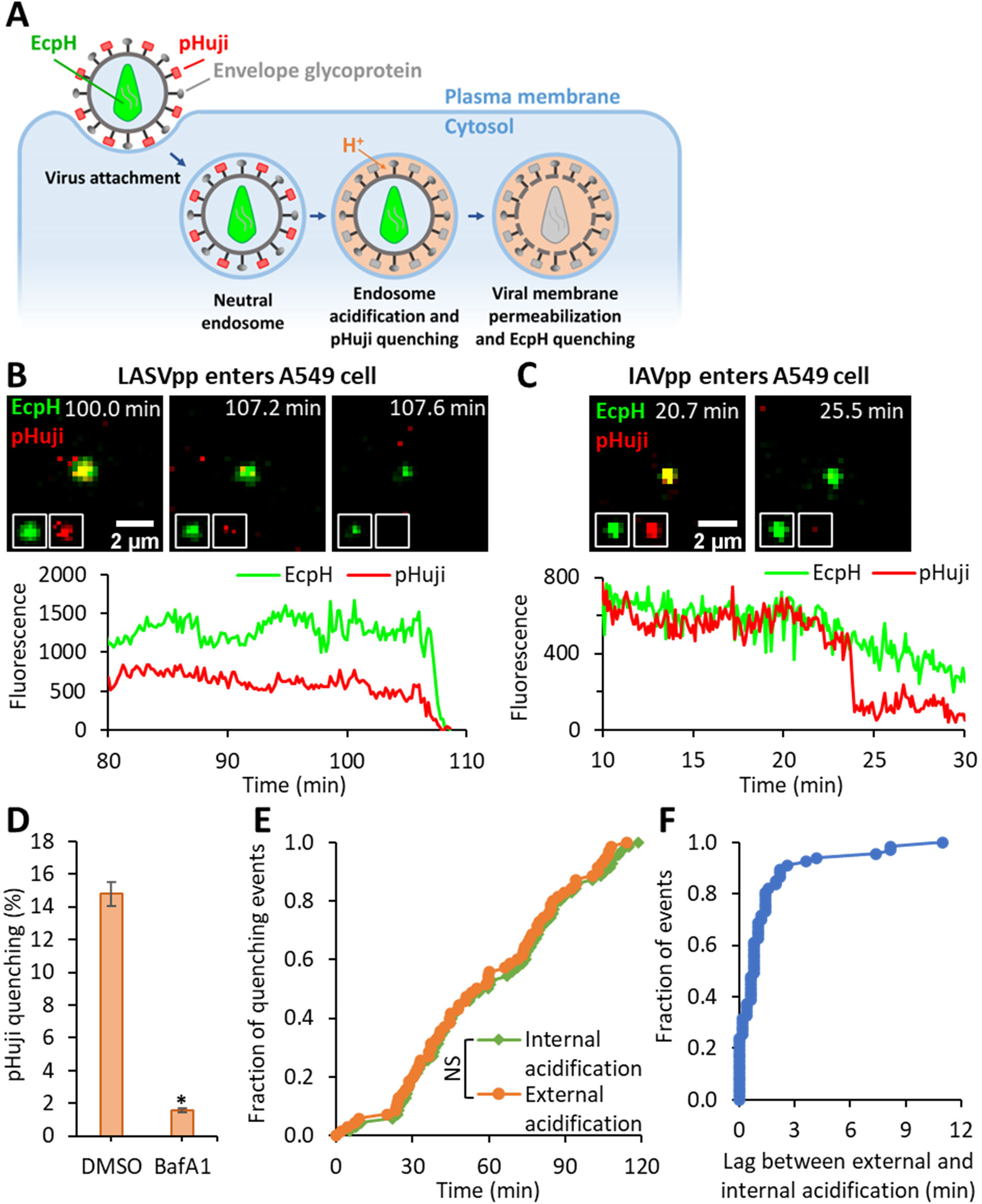
LASV pseudovirus interior acidification occurs shortly after entering acidic compartments. (A) Illustration of internalization and trafficking of LASVpp colabeled with pHuji and Gag-EcpH. LASVpp is internalized and trafficked to acidic endosomes where the viral surface probe, pHuji, is quenched. The LASVpp membrane permeability increases in acidic compartments, leading to the viral interior acidification and quenching of the internal low pH probe, EcpH. (B) Single LASVpp entry into acidic endosome and viral membrane permeabilization in A549 cells. Time-lapse images (top) and fluorescence traces (bottom) show that, shortly after virus entry into the acidic endosomes (pHuji quenching at 107.6 min), membrane permeabilization occurs, resulting in virus’ interior acidification (EcpH quenching) at 108.2 min (see Movie S4). (C) Single IAVpp entry into acidic endosome in A549 cells without membrane permeabilization. Time-lapse images (top) and fluorescence traces (bottom) show that IAVpp entry into the acidic endosome at 23.9 min leading to pHuji quenching with EcpH signal maintaining (see Movie S5). (D) Bafilomycin A1 (BafA1) inhibits pHuji quenching. Data shown are means ± SD of 2 independent experiments. Results were analyzed by Student’s t-test. *, p<0.05. (E) Kinetics of single LASVpp exterior and interior acidification (pHuji and EcpH quenching, respectively). Results were analyzed by Student’s t-test. NS, not significant. (F) Distribution of lag times between pHuji and EcpH quenching for each single pseudovirus.

Together, our results show that the LASVpp membrane is permeabilized shortly after entering acidified endosomes. Such a short lag time to membrane permeabilization and high pKa values of pHuji (∼7.7) and EcpH (∼7.1) [38, 39] suggest that virus membrane permeabilization occurs in early endosomes, soon after these compartments become mildly acidic.

### Cellular contact facilitates viral membrane permeabilization at low pH

To determine whether GPc interaction with cellular factors is required for viral membrane permeabilization at low pH, LASVpp labeled with mCherry-CL-YFP-Vpr were attached to coverslips, or to target cells in the cold to prevent virus uptake and fusion. Cell surface- or coverslip-bound viruses were then exposed to a membrane-impermeable pH 5.0 citrate buffer at 37 °C (Fig. 4A, B). Analysis of loss of the total YFP-Vpr fluorescence for all mCherry-labeled particles in the image field revealed an exponentially decaying signal that was markedly accelerated for cell-bound viruses compared to those attached to coverslips (Fig. 4C and Suppl. Fig. S6). The rate of the virion acidification was independent of the target cell type, displaying similar YFP decay constants in A549, HeLa and DF-1 cells (0.011, 0.017 and 0.013 sec^-1^, respectively, Fig. 4C). By contrast, a nearly instant quenching of YFP signal from 100% of particles was observed when a membrane-permeable acetic acid-based buffer was applied to viruses (Fig. 4C), supporting the existence of significant barrier for proton permeation through the viral membrane under basal conditions.

**Figure 4.**
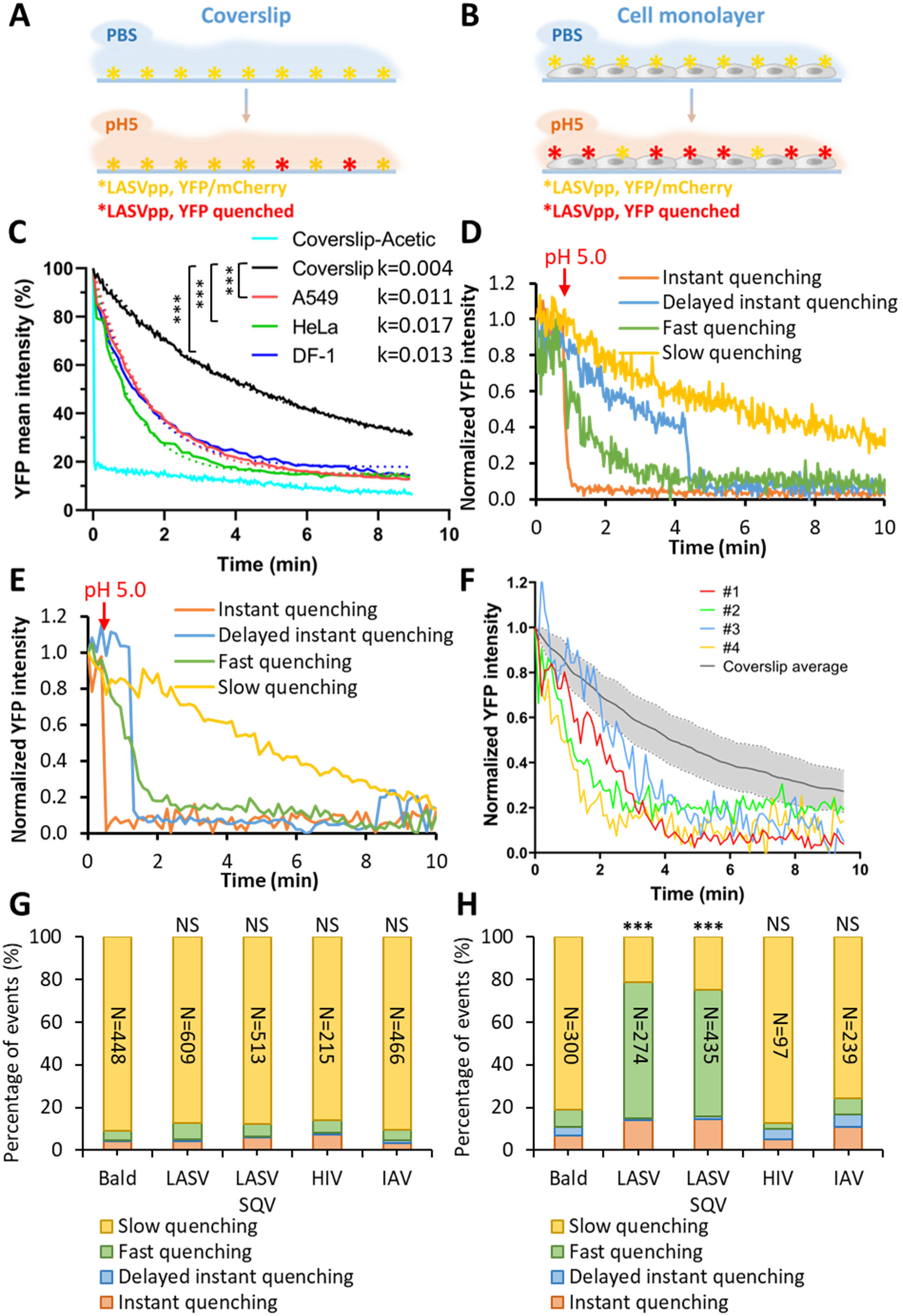
Virus-cell contact promotes LASVpp membrane permeabilization at low pH. (**A, B**) Illustration of the effects of LASVpp exposure to low pH on a coverslip (A) and on the cell surface (B). LASVpp were bound to the cell surface or to poly-L-lysine coated coverslips at 4 °C. GPc conformational changes are triggered by applying membrane impermeable pH 5.0 citrate buffer. (C) Mean YFP intensity decay of all coverslip-attached particles in the image field or on cells. Dotted-lines are single- exponential decay fits of data shown by solid lines. Control YFP quenching profile upon application of a membrane-permeable pH 5.0 acetic buffer is shown. Exponential decay rates k are in 1/sec. Results were analyzed by Student’s t test, ***, p<0.001; NS, p>0.05. (D, E) Representative examples of four types of YFP quenching of single LASVpp on coverslip (D) or on cells (E). The point of adding a low pH buffer is marked with the red arrows. (F) Examples of the fast single virus YFP quenching events in A549 cell surface (#1-#4) overlaid onto the average YFP intensity profile for single LASVpp acidified on coverslip. The shaded area represents standard deviation of the mean YFP decay curve. (G, H) Quantification of the above four categories of YFP-Vpr quenching of single LASVpp on coverslip (G) or on DF-1 cells (H). Numbers within the bars are the total numbers of particles analyzed by Fisher’s exact test. ***, p<0.001; NS, not significant.

To gain further insights into the loss of total YFP signal from multiple viral particles at low pH (Fig. 4C), we tracked the YFP-Vpr signal of single LASVpp on coverslips or on the cell surface upon shifting to pH 5.0. Single virus fluorescence traces exhibited distinct YFP quenching profiles and were classified into 4 categories (Fig. 4D-F): (1) “instant” quenching upon acidification manifested by an abrupt drop of YFP signal to background level for a small fraction (3.9%, Suppl. Fig. S4) of particles with preexisting membrane defects; (2) “delayed instant” quenching after an initial slow decay of fluorescence, also in a very small fraction of particles, suggesting a variable lag to rapid membrane permeabilization; (3) “slow” quenching, likely resulting from the baseline proton permeability of the viral membrane; and (4) “fast” gradual quenching that appears to occur due to an increase in the baseline viral membrane permeability. (Additional examples of these distinct YFP-Vpr quenching events on coverslips and cells are shown in Suppl. Fig. S7). Single YFP quenching events were considered fast, if the YFP intensity fell below a standard deviation from the average slow YFP quenching curve for viruses on coverslips (Fig. 4F). It should be noted that, judging by the gradual loss of YFP signal over the course of minutes, fast YFP quenching events appear to report a rather moderate membrane permeability increase. From the kinetics of fast YFP quenching in single virions, we estimated the membrane permeability to be on the order of 4ꞏ10^-8^ cm/sec, which is only an order of magnitude higher that the estimated permeability corresponding to slow YFP quenching events (Suppl. Fig. S8).

Approximately 88% of coverslip-adhered LASVpp exhibited slow/baseline YFP-Vpr quenching (Fig. 4D, G), likely reflecting background proton permeability of the viral membrane; only ∼7.2% exhibited fast YFP quenching after low pH application. In sharp contrast, cell-bound viruses exhibited a dramatic increase in the fraction of fast YFP quenching events (∼60%) and only ∼20% exhibited slow/baseline quenching (Fig. 4H). Other YFP quenching categories (instant and delayed instant) did not exhibit significant changes between cell-free and cell-attached particles.

Our results imply that efficient virion membrane permeabilization requires conditions that are permissive for GPc-mediated membrane fusion: low pH and virus contact with cells. This notion is supported by findings that “bald” particles lacking any viral glycoprotein or pseudoviruses expressing HIV-1 Env or IAV hemagglutinin (HA) do not exhibit fast YFP quenching on coverslips or when attached to cells (Fig. 4G, H). The invariant fraction of fast YFP quenching events for coverslip-attached and cell-bound HIV-1 Env and IAV HA pseudotyped particles was not due to poor incorporation of these glycoproteins into HIV-1 particles (Suppl. Fig. S9). In contrast, all control viruses exhibit predominant slow quenching, supporting the notion that the slow YFP quenching rate for these viruses likely represents passive (baseline) proton permeability of the viral membrane. We therefore hypothesized that fast YFP quenching is likely mediated by LASV GPc. We also tested if maturation of the HIV-1 core, which is known to regulate the Env function [40–42], affects viral membrane permeability. Immature LASVpp produced in the presence of the HIV-1 protease inhibitor, Saquinavir (SQV), exhibited acid-induced permeability increases on cells and not on coverslips, similar to the changes observed for mature particles (Fig. 4G, H). Thus, HIV-1 maturation does not affect the ability of GPc to compromise the membrane’s barrier function.

Together, the above results show that virus-cell contact at low pH is needed for efficient viral membrane permeabilization by LASV GPc. To determine whether specific LASV GPc interactions with the cellular receptors, α-dystroglycan or LAMP1, can promote the increases in viral membrane permeability, coverslip-attached pseudoviruses were pretreated with recombinant α-dystroglycan or a soluble fragment of LAMP1 (sLAMP1) and exposed to a low pH buffer supplemented with these proteins. Under these conditions, α-dystroglycan or sLAMP1 did not accelerate the loss of YFP fluorescence at low pH (Suppl. Fig. S10). Since α- dystroglycan has been shown to dissociate from GPc at low pH [18], its failure to modulate the viral membrane permeability under in these experiments is perhaps not surprising. The lack of a detectable effect of soluble LAMP1 may indicate the need for membrane anchored LAMP1 to support GPc conformational changes associated with the virion membrane permeabilization.

Evidence indeed suggests that membrane cholesterol may be required for functional LASV GPc interaction with LAMP1 [43].

### The increase in LASVpp membrane permeability is caused by fusion-inducing conformational changes in GPc

Having established that virion membrane permeabilization is mediated by LASV GPc upon exposure to low pH and contact with the target cell membrane, we next examined the link between the membrane permeability increases and virus fusion. Our results show that only ∼14% of all A549 cell-bound LASVpp are internalized into acidic compartments (Fig. 3D) and that ∼30% of these particles fuse within 2 hrs of infection (Suppl. Fig. S2 and [31, 32]). We tracked randomly selected single-pseudovirus YFP-Vpr quenching events for 2 hrs and categorized these depending on whether particles did or did not undergo membrane fusion (as judged by release of mCherry). We plotted the ensemble average profiles of single particle YFP-Vpr quenching events (aligned to the onset of quenching) and determined the decay constants for fusing and non-fusing LASVpp (Fig 5A, B). Both the average profiles and individual rate constants of fusing particles showed a highly significant enhancement of membrane permeabilization relative to non-fusing particles. Approximately 80% of fusing LASVpp exhibited fast YFP-Vpr quenching, in contrast to only ∼20% in non-fusing particles (Fig. 5C). Consistent with fusion-related increases in LASVpp membrane permeability, non-fusing particles exhibited similar YFP quenching profiles to particles attached to coverslips, with no detectable changes in the baseline proton permeability (Fig. 5A, B).

**Figure 5.**
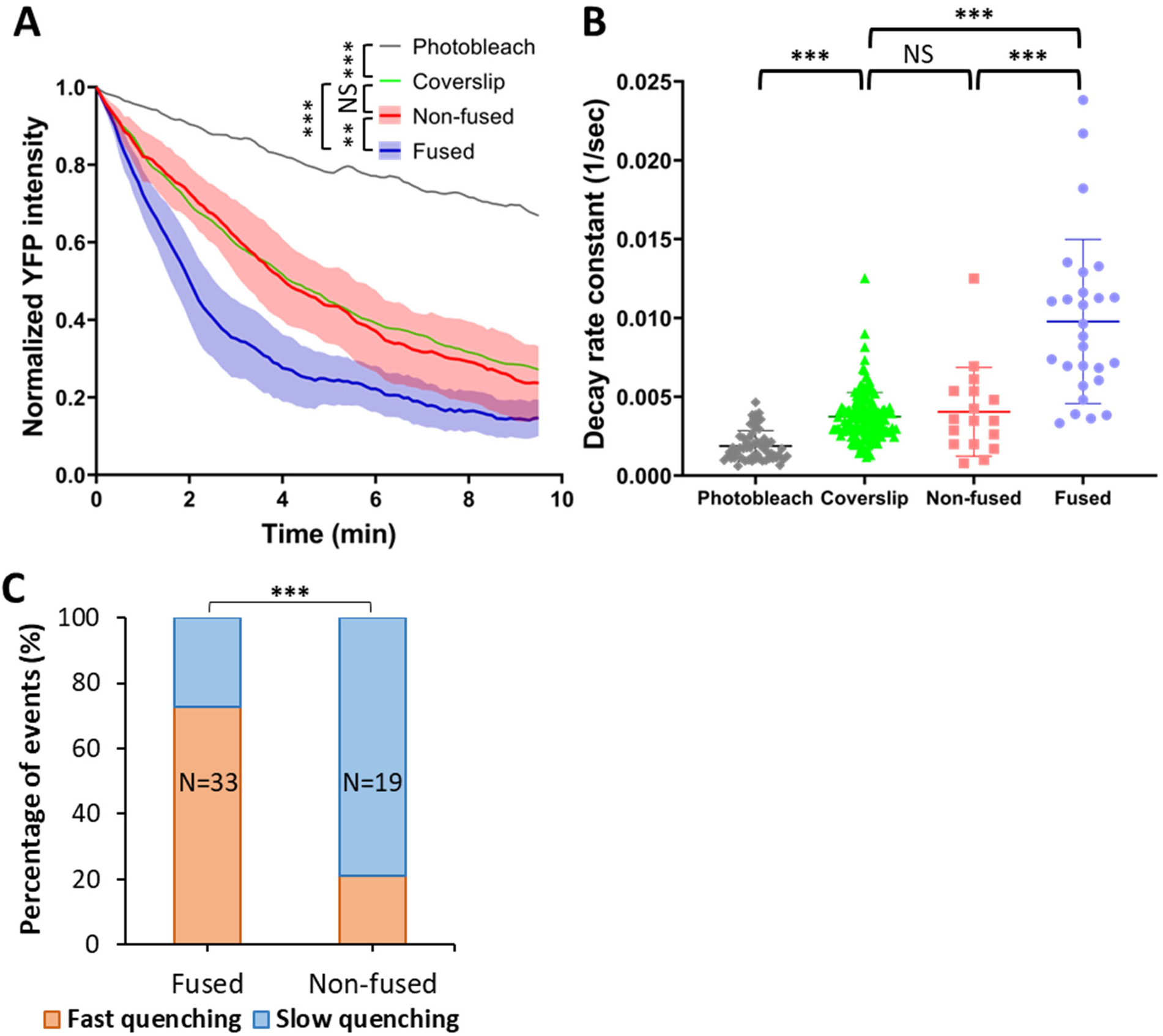
Membrane permeability increases are observed for fusing but not non-fusing LASVpp. (A) Normalized average single YFP-Vpr intensity profiles of LASVpp upon exposure to low pH on a coverslip or upon entry into A549 cells through a conventional endocytic pathway (fused vs. non-fused particles). Individual YFP intensity decays are aligned at the onset of quenching. Photobleaching-mediated YFP signal decay in PBS^+/+^ is shown as reference. Shadowed area is the 95% confidence interval. Data were analyzed by two-way ANOVA. **, p<0.01; ***, p<0.001; NS, not significant. (B) The YFP-Vpr intensity decay rates for fusing vs non-fusing particles, as well as particles on a coverslip obtained by single-exponential fitting are shown. The rate of YFP photobleaching is also shown. Data were analyzed by Student’s t-test. ***, p<0.001. (C) Quantification of different types of YFP-Vpr quenching of fused or non-fused single LASVpp in live cell analyzed by Fisher’s exact test.

To further investigate the relationship between the fusogenic conformational changes in GPc and virion membrane permeabilization, we examined a series of GPc mutants that are defective in fusion, as well as the effects of a small-molecule fusion inhibitor. For these studies, we utilized a virus-cell fusion assay, which is based on the cytosolic delivery of the pseudovirus-incorporated β-lactamase-Vpr construct [44, 45]. The K33A mutation lies within SSP, in a region that appears to be involved in pH sensing and the activation of membrane fusion [46, 47]. The fusion-defective K33A GPc was efficiently incorporated into pseudovirions (see Suppl. Fig. S11), which displayed the expected inability to induce fusion [47]. LASVpp bearing this mutant produced a near background level of fusion signal (Fig. 6A). Parallel imaging experiments revealed impaired ability of the K33A mutant to increase membrane permeability of cell-attached pseudovirions upon exposure to low pH (Fig. 6B). Accordingly, the fraction of the fast single-virus YFP quenching events was greatly reduced from ∼80% for wild-type GPc to 10% for the K33A mutant (Fig. 6C).

**Figure 6.**
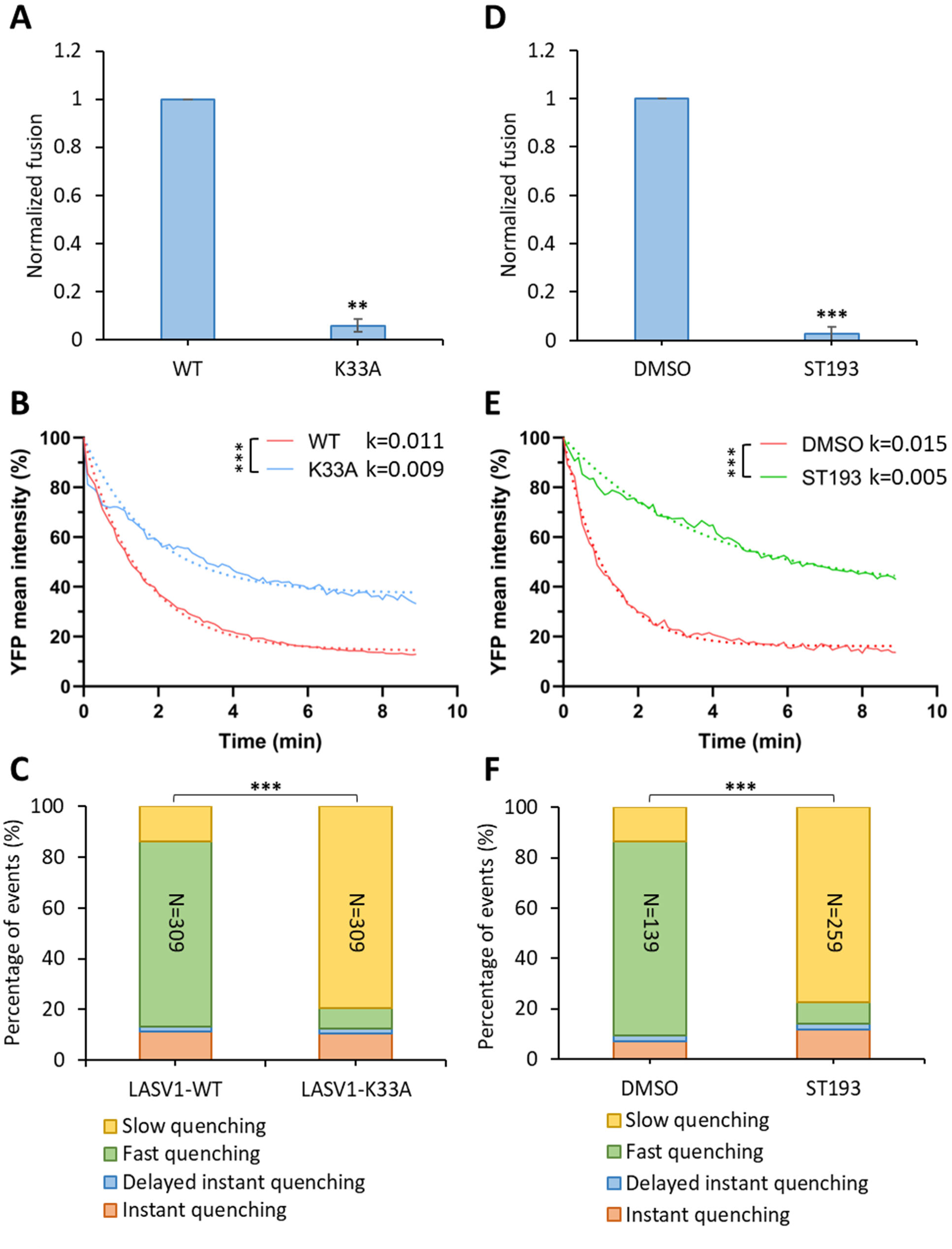
Fusion-impairing LASV GPc mutation and fusion inhibitor impair viral membrane permeabilization. (A) Wild-type and K33A mutant LASVpp-BlaM fusion with A549 cells. LASVpp was bound to A549 cells in the cold and viral fusion was initiated by shifting to 37 °C and incubating for 2 hours. Data shown are means ± SD of 3 independent experiments. Results were analyzed by Student’s t-test. **, p<0.01. (B) Mean YFP-Vpr intensity decay of LASV GPc WT and K33A mutant on the surface of A549 cells after applying membrane-impermeable pH 5.0 citrate buffer. K33A mutant abrogate LASVpp fusion. The exponential decay rates k are in 1/sec. Results were analyzed by Student’s t-test, ***, p<0.001. (C) Quantification of different types of single LASVpp YFP-Vpr quenching events for LASVpp WT or K33A GPc mutant on A549 cells after applying low pH. (D) ST-193 inhibits LASVpp- BlaM fusion with A549 cells. LASVpp was bound to A549 cells in the cold and viral fusion was initiated by shifting to 37 °C and incubating for 2 hours in the presence of 10 µM ST-193 or equal volume of solvent (DMSO). Data shown are means ± SD of 3 independent experiments. Results were analyzed by Student’s t-test. ***, p<0.001. (E) Mean YFP-Vpr intensity decay of LASVpp on A549 cell surface after applying pH 5.0 citrate buffer in the presence or absence of 10 µM of ST-193. Exponential decay rates k are shown in 1/sec. Results were analyzed by Student’s t test, ***, p<0.001. Note that the different rates of YFP quenching for WT GPc in panels B and E are due to the presence of DMSO (vehicle) in experiments with ST-193. (F) Quantification of different types of single LASVpp YFP-Vpr quenching events on A549 cells in the presence or absence of 10 µM ST-193, after applying low pH. Data in panels (C) and (F) were analyzed by Fisher’s exact test. ***, p<0.001.

We also tested the effect of small molecule LASVpp fusion inhibitor, ST-193, which is thought to prevent early GPc conformational changes induced by low pH [48, 49]. ST-193 abrogated LASVpp fusion with A549 cells (Fig. 6D) and inhibited the decay of average YFP-Vpr signal at low pH (Fig. 6E). Accordingly, single particle tracking demonstrated a greatly reduced fraction of fast YFP-Vpr quenching events for ST-193-treated viruses compared to an untreated control (Fig. 6F). A correlation between LASVpp fusion activity and fast YFP-Vpr quenching implies that the latter events are caused by on-path conformational changes in GPc. Again, slow YFP quenching events likely reflect background proton permeability. Together, these results suggest that membrane permeabilization is intrinsic to events in GPc-induced fusion, and again point to the role of GPc conformational changes in mediating this change in membrane permeability.

To extend these studies, we tested a panel of additional GPc mutants that have been shown to variously impair fusion activity. The H230Y and H230E mutations in GP1 render GPc defective in LAMP1 binding, reducing LASV infectivity but not GPc-mediated syncytia formation [17, 50]. The D268A, L270A, R282A and I323A mutations target different domains in GP2 (fusion-peptide proximal, internal fusion loop, and heptad repeat domain 1, respectively) and each attenuates GPc fusion activity [51]. These mutants are incorporating into pseudoviruses at levels comparable to wild-type GPc (Suppl. Fig. S11) and exhibit strongly reduced fusion activity, as measured by a β-lactamase-Vpr based assay (Fig. 7A). The only exception was the H230E mutant, which was incorporated at a lower level compared to the wild-type glycoprotein.

**Figure 7.**
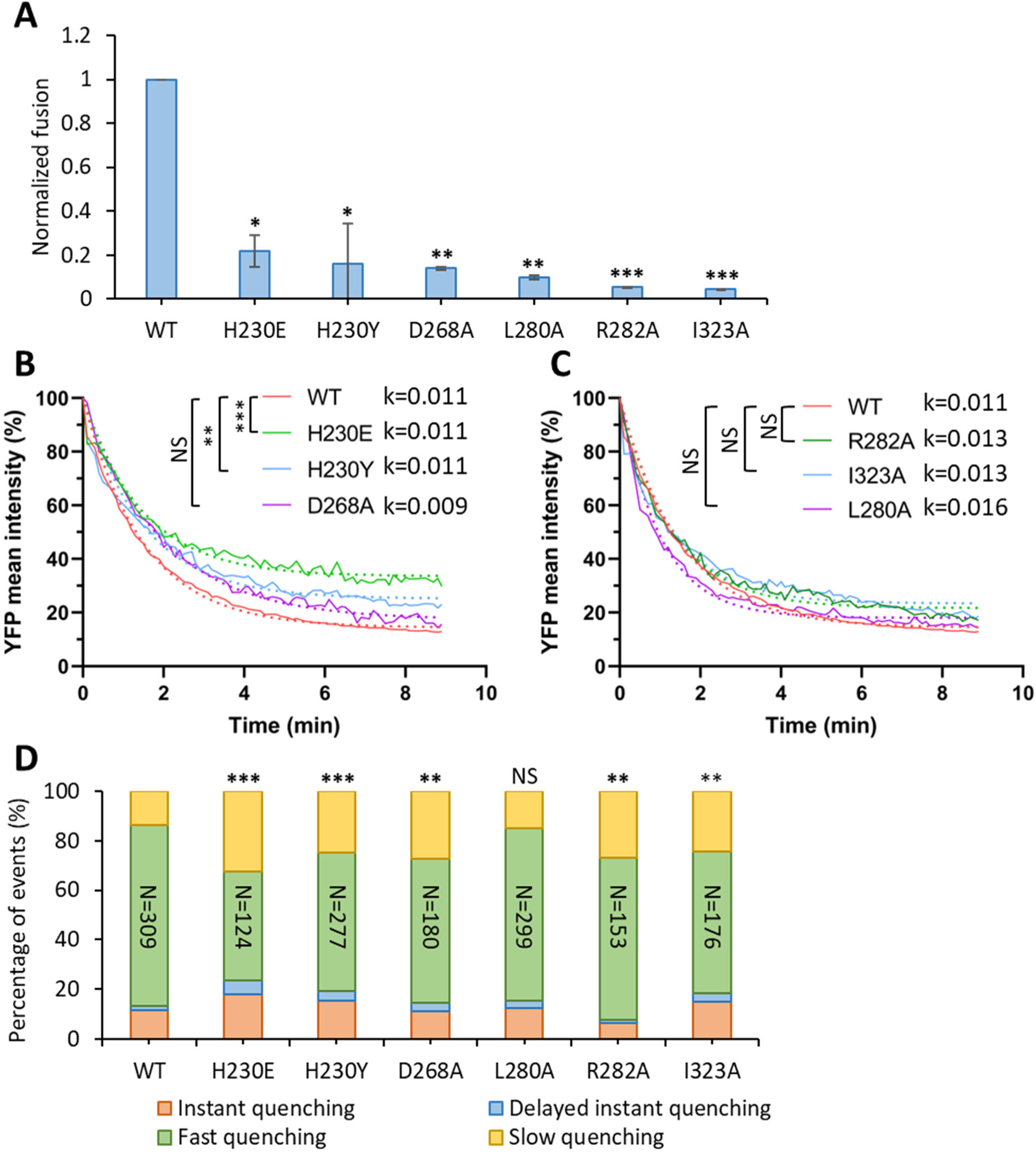
LASVpp membrane permeabilization is inhibited by fusion attenuating GPc mutations. (A) Wild-type and mutant LASVpp fusion with A549 cells measured by a BlaM assay. LASVpp was bound to A549 cells in the cold, and the fusion was initiated by shifting to 37 °C and incubating for 2 hours. Data shown are means ± SD of 3 independent experiments. Results were analyzed by Student’s t-test. *, p<0.05; **, p<0.01; ***, p<0.001. (B, C) Mean YFP-Vpr intensity decay of LASVpp mutants on A549 cell surface after applying membrane- impermeable pH 5.0 citrate buffer. The exponential decay rates k are shown in 1/sec. Results were analyzed by Student’s t-test, **, p<0.01; ***, p<0.001; NS, not significant. (D) Quantification of different types of YFP-Vpr quenching of single LASVpp GPc mutants on A549 cells after applying low pH. Data was analyzed by Fisher’s exact test. **, p<0.01; ***, p<0.001; NS, not significant.

Interestingly, only the two H230 mutations in the GP1 subunit significantly reduced the rate of YFP quenching upon exposure of cell-bound viruses to low pH (Fig. 7B). By contrast, neither of the GP2 mutations tested, significantly slowed down YFP quenching (Fig. 7C). In agreement with these results, the fraction of accelerated YFP quenching events was significantly reduced by the two H230 mutations, whereas the GP2 mutants displayed either a more moderate effect or had no significant effect (Fig. 7D). Since the relatively minor fraction of instant YFP quenching events (Fig. 4) does not considerably vary across the mutants (in the 8-23% range), these events are unlikely to be associated with GPc refolding. Collectively, these results highlight the relationship between membrane permeabilization and GPc function in membrane fusion, but do not allow us to define a specific point in the fusion process at which permeabilization is activated.

## Discussion

Viral protein-mediated membrane fusion has been extensively studied for many viruses and using a variety of experimental systems. Despite structural diversity, leading to classification of viral glycoproteins into three distinct classes [52–54], the general mechanism by which these proteins induce membrane fusion appears to be conserved. The predominant model posits that, upon triggering, viral glycoproteins undergo large-scale conformational changes involving at least two critical steps – exposure of the hydrophobic fusion peptide domain and insertion into the target membrane, followed by refolding into a hairpin-like structure that brings the fusion peptides and transmembrane domains into proximity. This latter step is thought to overcome the activation energy barrier for membrane fusion. The lipid rearrangements *en route* to membrane merger also appear to be conserved. It is widely accepted that protein-mediated membrane fusion progresses through a hemifusion intermediate, whereby the contacting outer leaflets of the virus and cell membranes are merged, but the inner leaflets remain distinct and form a hemifusion diaphragm [55–57]. Progression through a hemifusion structure is thought to ensure “tight” fusion that prevents leakage of the cell/viral content into the external milieu. In agreement with the “tight” fusion model, our methodologies routinely find that membrane fusion by many viral glycoproteins (IAV HA, VSV G and ALSV Env) occurs without compromising the barrier function of the viral membrane, i.e., we detect no quenching of an intraviral pH-sensor in acidic endosomes prior to virus fusion [21–31] (but see [58–60] for reports of “leaky” fusion by influenza HA). In contrast, single pseudovirus fusion mediated by LASV GPc occurs after permeabilization of the viral membrane [31, 32].

Our results strongly suggest that arenaviral membrane permeabilization is mediated by fusion-inducing conformational changes in GPc. First, glycoproteins of other viruses, such as HIV-1 or IAV, do not cause membrane permeability increases at low pH. Second, an increase in membrane permeability at low pH is selectively observed for LASVpp that successfully fuse and rarely for non-fusing particles. Third, mutations that impair LASV fusion activity or small-molecule fusion inhibitors both interfere with the GPc’s ability to permeabilize the virus membrane at low pH. Fourth, virus-cell contact greatly augments the viral membrane permeability increase at low pH, as compared to cell-free viruses. The failure of soluble LAMP1 to promote cell-free virus permeabilization at low pH appears to highlight the importance of LASV binding to LAMP1 in the context of a target membrane for driving on-path GPc refolding and viral membrane permeabilization. Indeed, the GPc-LAMP1 interaction has been reported to depend on cholesterol [43]. We further demonstrate that the membrane permeability increase is observed in Old- and New-World arenavirus GPcs, in both HIV-1 pseudovirions and arenavirus VLPs, and is independent of target cell types. Together, these results imply that functional conformational changes in GPc triggered by endosomal acidification, and interactions with unknown host factors, are required for the permeability increases in the viral membrane prior to the completion of membrane fusion.

The observation that YFP-Vpr quenching occurs shortly after the initial acidification of endosomal lumen (Fig. 3E) indicates that conformational changes in LASV GPc responsible for virion membrane permeabilization occur in early endosomes. Since LASV is thought to fuse with LAMP-1-conaining late endosomes/endolysosomes [8, 18, 19, 61], it is possible that permeabilized viruses are trafficked to and fuse with these compartments. Indeed, the average lag time between YFP-Vpr quenching and nascent LASVpp fusion pore formation in DF-1 cells expressing human LAMP1 was ∼8 min [32], which may be sufficient for virus transport to late compartments.

Analysis of different fusion-impaired mutants suggests that mutations interfering with early conformational changes of LASV GPc (e.g., K33A which is thought to prevent initiation of GPc refolding at low pH [47]) appear to more potently inhibit the viral membrane permeabilization than those affecting the late steps of fusion (e.g., I323A in the GP2 HR1 region [51] that is involved in the final trimeric hairpin formation). ST-193, which has been hypothesized to block early conformational changes in GPc by targeting the interface between the stable signal peptide (SSP) and the transmembrane domains of GP2 [49, 62], also potently inhibited the membrane-permeabilizing activity of this protein (Fig. 6D-F). We note, however, that our recent data indicate that ST-193 captures LASV GPc-mediated cell-cell fusion at a hemifusion stage [7]. Since hemifusion generally requires the fusion peptide insertion into the target membrane, this finding is more in line with the inhibitor blocking a late conformational change in GPc.

Fast YFP quenching, which correlates with GPc’s fusion activity, leads to gradual acidification of the virion interior, consistent with a modest increase in membrane proton permeability. It is thus possible that subtle structural changes at the interface between the SSP and GP2 transmembrane domain, which are antagonized by ST-193, may cause the observed increases in the viral membrane permeability. It is also possible that conformational changes in GPc create lipid packing defects that result in a moderately increased proton permeability. Irrespective of the mechanism, increased virion membrane permeability is associated with the fusogenic structural changes in GPc suggesting that acidification of the virus interior may play a role virus entry.

Indeed, a recent report has shown that low pH promotes Z protein oligomerization [63], suggesting that the virion interior acidification can alter the matrix protein arrangement and prime the viral membrane for fusion. Moreover, the same study has also shown that Z and NP interaction is disrupted by acidic pH. It is thus likely that the arenavirus GPc-mediated acidification of the virion interior plays a role in post-fusion steps of infection, including viral ribonucleoprotein release into the cytosol. This notion is akin to the role of the M2 viroporin of the Influenza virus, which has been found to be essential for uncoating of the nucleocapsid [64, 65]. Future experiments may shed light on the molecular mechanism underlying the membrane-permeabilizing activity of arenavirus GPc and its role in the early steps of arenavirus infection.

## Materials and Methods

### Plasmids, cell lines and reagents

The pR9ΔEnv, mCherry-2xCL-YFP-Vpr, pMM310 BlaM-Vpr, pcRev, EcpH-ICAM, pCAGGS- WSN-HA, pCAGGS-WSN-NA have been described previously [20, 22, 66, 67]. The LASV-GPc plasmid was a gift from Dr. François-Loic Cosset (Université de Lyon, France) [68]. LASV-GPc-K33A, JUNV-GPc, JUNV GPc, NP and Z (all Candid-1 strain) were constructed by Jack Nunberg (unpublished results). The PolI-S Can reverse genetics S-RNA of Candid-1 JUNV plasmid was a gift from Dr. Juan Carlos de la Torre (The Scripps Research Institute) [69].

LASV-GPc-D268A, LASV-GPc-L280A, LASV-GPc-R282A and LASV-GPc-I323A were described previously [51]. MACV-GPc was a gift from Dr. Michael Farzan (Scripps Research Institute) [70]. LCMV-GPc was a gift from Shan-Lu Liu (The Ohio State University). pCAGGS- HXB2 was provided by Dr. James Binley (Torrey Pines Institute, CA) [71]. pDisplay-pHuji was from Addgene (Cat# 61556). Gag-EcpH was constructed by Dr. Kosuke Miyauchi. EcpH was amplified by PCR and replaced the mCherry in pcDNA3.1zeo+HIV Gag-mCherry by *BamHI/XhoI* restriction enzyme digestion and ligation.

To label the JUNV-NP (Candid-1 stain), YFP was inserted at an exposed loop by replacing the D93 residue with the YFP sequence [36]. The inserted YFP lacks the initiation and termination codons and is flanked by GSSG linkers. The YFP sequence was amplified by PCR (forward primer: 5’ CCA TGA GGA GTG TTC AAC GAA ACA CAG TTT TCA AGG TGG GAA GCT CCG GCG TGA GCA AGG GCG AGG AGC TG; reverse primer: 5’ GGT CAG ACG CCA ACT CCA TCA GTT CAT CCC TCC CCA GGC CGG AGC TGC CCT TGT ACA GCT CGT CCA TGC CGA G) and the resulting megaprimer was inserted at the D93 site (replacing D93 residue) in the PolI-S Can plasmid by QuikChange mutagenesis (Agilent Technologies, Santa Clara, CA). NP-DYFP then was adapted and extracted from the modified PolI-S Can plasmid by PCR (forward primer: 5’ CGC GGC TAG CTC TGG CAT GGC ACA CTC CAA GGA GGT TCC AAG C; reverse primer: 5’ CGC GCT CGA GTG CTT ACA GTG CAT AGG CTG CCT TCG G) and inserted into pcDNA3.1(+) by *NheI/XhoI* restriction enzyme digestion and ligation. The NP-DYFP insert was then transferred to the plasmid pCAGGS by *SacI/XhoI* restriction enzyme digestion and ligation.

To obtain JUNV-NP-DmCherry (Candid-1 strain), mCherry sequence was amplified by PCR (forward primer: 5’ CCA TGA GGA GTG TTC AAC GAA ACA CAG TTT TCA AGG TGG GAA GCT CCG GCA TGG TGA GCA AGG GTG AGG AGG; reverse primer: 5’GGT CAG ACG CCA ACT CCA TCA GTT CAT CCC TCC CCA GGC CGG AGC TGC CCT TGT ACA GCT CGT CCA TGC CGC C) and inserted and replaced YFP in pCDNA3.1(+)-JUNV-NP- DYFP by QuickChange PCR with the mCherry segment served as megaprimers. The NP- DmCherry insert was then transferred to the plasmid pCAGGS by SacI/XhoI restriction enzyme digestion and ligation.

Human embryonic kidney HEK293T/17 cells, human lung epithelial A549 cells, human cervix epithelial HeLa cells, human bone U2OS cells, African green monkey kidney VeroE6 and fibroblast CV-1 cells, as well as chicken embryonic fibroblast DF-1 cells, were obtained from ATCC (Manassas, VA, USA). All cells were grown in high glucose Dulbecco’s Modified Eagle Medium (DMEM; Mediatech, Manassas, VA, USA) containing 10% heat-inactivated Fetal Bovine Serum (FBS; Atlanta Biologicals, Flowery Branch, GA, USA) and 1% penicillin/streptomycin (GeminiBio, West Sacramento, CA, USA). The growth medium for HEK 293T/17 cells was supplemented with 0.5 mg/ml G418 (Genesee Scientific, San Diego, CA, USA).

ST-193 was purchased from MedChemExpress (NJ, USA). BSA and Bafilomycin A1 were acquired from Sigma-Aldrich (MO, USA). The recombinant human α-dystroglycan (rhDG) protein was acquired from R&D Systems (MN, USA). The LAMP1 (H4A3) mouse antibody (LAMP1ab) was purchased from Santa Cruz Biotechnology (TX, USA).

### Pseudovirus and virus like particle production

The pseudoviruses and virus like particles (VLPs) were produced by transfecting HEK293T/17 cells using JetPRIME transfection reagent (Polyplus-transfection, Illkirch-Graffenstaden, France). HEK293T/17 cells were seeded in DMEM with 10% FBS the day before transfection. To produce the pseudoviruses co-labeled with YFP and mCherry for single virus fusion experiments, HEK293T/17 cells were transfected with pR9ΔEnv, mCherry-CL-YFP-Vpr, pcRev and different envelope glycoprotein plasmids: LASV-GPc wild-tpe or mutants, MACV-GPc, JUNV-GPc, LCMV-GPc, HIV-HXB2 Env and IAV WSN HA and NA. These viral envelope encoding plasmids were omitted when producing “bald” control pseudoviruses. To produce immature LASV pseudovirus (LASV-SQV), HEK293T/17 cells were transfected with the above mixture of plasmids and maintained in the presence of 300 nM of the HIV protease inhibitor Saquinavir (SQV, contributed by DAIDS/NIAID, NIH HIV Reagent Program). To produce LASV and IAV pseudoviruses co-labeled with the surface and core pH-sensors for single virus internalization experiments, HEK293T/17 cells were transfected with LASV-GPc or IAV WSN HA and NA, pR9ΔEnv, pDisplay-pHuji, HIV-1 Gag-EcpH, and pcRev.

To produce VLPs with YFP and mCherry labeling for single VLP fusion experiments, HEK293T/17 cells were transfected with plasmids encoding the following Candid-1 strain of JUNV-derived constructs: NP-DYFP, NP-DmCherry, Z and GPc glycoproteins of (LASV, JUNV, Candid-1, and MACV. To produce LASV pseudoviruses carrying β-lactamase-Vpr (BlaM-Vpr) for a bulk virus-cell fusion assay [44, 45]. HEK293T/17 cells were transfected with wild-type or mutant LASV-GPc, pR9ΔEnv, BlaM-Vpr and pcRev plasmids. Supernatants were collected at 48 hours post-transfection and filtered through a 0.45 µm syringe filter to remove cell debris and virus aggregates. LASVpp-BlaM viruses were concentrated ten times with Lenti-X concentrator (Clontech Laboratories, Mountain View, CA, USA), snap-frozen and stored at - 80 °C.

### Immunostaining for envelope glycoproteins

Viruses were incubated with poly-L-lysine coated 8-well chambered glass coverslip at room temperature for 30 minutes, washed with PBS^+/+^, and fixed with 4% paraformaldehyde (Electron Microscopy Sciences, PA, USA) at room temperature for 15 min. Paraformaldehyde was quenched by washing 5 times with 20 mM Tris in PBS^+/+^. Samples were then blocked with 15% FBS in PBS^+/+^ at room temperature for 2 hours. To detect LASV-GPc, LASV pseudoviruses were incubated with 10 µg/ml of 12.1F human anti-LASV-GPc primary antibody (Zalgen Labs, MD, USA) [72] at 4 °C overnight followed by staining with 2 µg/ml of secondary AlexaFluor647-conjugated Donkey anti-human IgG antibody (ThermoFisher Scientific, OR, USA) at room temperature for 45 min. To detect HIV-1 HXB2 Env, pseudoviruses were incubated with 5 µg/ml of 2G12 human anti-Env primary antibody at 4 °C overnight and 2 µg/ml AlexaFluor647 donkey anti-Human IgG secondary antibody at room temperature for 45 min. To detect IAV HA, pseudoviruses were incubated with Rabbit anti-IAV WSN HA R2376 polyclonal antibody (1:100 diluted, a gift of Dr. David Steinhauer) at 4 °C overnight and 2 µg/ml AlexaFluor647 Donkey anti-rabbit IgG secondary antibody (ThermoFisher Scientific Corporation, OR, USA) at room temperature for 45 min. Samples were washed 5 times with 15% FBS/PBS^+/+^ after incubation with both primary and secondary antibodies. Images were acquired on a DeltaVision microscope using Olympus 60x UPlanFluo /1.3 NA oil objective (Olympus, Japan). Co-localization and immunofluorescence were analyzed with the ComDet plugin in ImageJ.

### Single virus and VLP imaging in live cell

Target cells were seeded the day before imaging in 35 mm collagen coated glass-bottom Petri dishes (MatTek, MA, USA) in Fluorobrite DMEM (Life Technologies Corporation, NY, USA) containing 10% FBS, penicillin, streptomycin, and L-glutamine. Sixty µl of pseudoviruses or VLPs diluted with PBS^+/+^ (30-120-fold, depending on the virus’ concentration) were bound to the cells by centrifugation at 1550xg for 20 min, 4 °C. Unbound viruses or VLPs were removed by washing with cold PBS^+/+^. Virus or VLP entry and fusion were initiated by adding 2 ml of warm Fluorobrite DMEM supplemented with 10% FBS and 20 mM HEPES (pH 7.2). Samples were placed on a DeltaVision Elite microscope equipped with a temperature, humidity and CO2- controlled chamber and immediately imaged in a time-lapse mode for 2 h. Every 6 sec, 4 Z-stacks spaced by 1.5 μm were acquired to span the thickness of cells using Olympus 60x UPlanFluo /1.3 NA oil objective (Olympus, Japan).

For particle tracking and analyses, 3D images were converted to maximum intensity projections. Single particle fusion or pH-sensitive probe quenching events were annotated with the ROI manager tool in ImageJ. Single particle tracking was performed using ICY image analysis software (icy.bioimageanalysis.org). Briefly, pseudoviruses or VLPs were identified by the Spot Detection plugin and tracked using the Spot Tracking plugin to determine their trajectories and changes in fluorescence intensity over time.

### Single virus *in vitro* acidification

To assess changes in membrane permeability resulting in acidification of virus’ interior, viruses were bound to poly-L-lysine coated 8-well chambered glass coverslip at 4 °C for 30 min, washed with cold PBS^+/+^ to remove unbound viruses and kept in 100 µl PBS^+/+^. Time-lapse images with a time interval of 2 seconds were acquired on a DeltaVision microscope using Olympus 60x UPlanFluo /1.3 NA oil objective at 37 °C for 12 min. To lower the pH, 700 µl of warm membrane-impermeable pH 5.0 citrate buffer was added to each well 20 seconds after the onset of imaging.

To monitor the membrane permeability of viruses attached to the cell surface, cells were seeded in 35 mm collagen coated glass-bottom Petri dishes in Fluorobrite DMEM containing 10% FBS, penicillin, streptomycin, and L-glutamine the day before imaging. Viruses were bound to the cells by centrifugation at 1550xg for 20 min, 4 °C. Unbound viruses were washed off with cold PBS^+/+^ and covered with 70 µl of PBS^+/+^. Time-lapse imaging was performed on a DeltaVision microscope for 12 min using Olympus 60x UPlanFluo /1.3 NA oil objective. Every 8 sec, 4 Z-stacks spaced by 1.5 μm were acquired to cover the thickness of cells. At 20 seconds after starting imaging, 500 µl of warm pH 5.0 citrate buffer was added to lower the external pH. Maximum intensity projections of 3D images were used for subsequent analysis.

To analyze the YFP mean intensity change over time in low pH buffer, particles were detected and their YFP intensity was measured by the ComDet plugin in ImageJ with mCherry as a reference color. The YFP mean intensity were calculated in Excel and fitted to a single-exponential decay model in GraphPad Prism.

To monitor membrane permeabilization and virus interior acidification at the single virus level, particles were tracked using the ICY software. Labeled pseudoviruses were identified by Spot Detection plugin and tracked using Spot Tracking plugin to determine the fluorescence intensity over time. Based on the single particle intensity change, YFP quenching was divided into four types: instant quenching, delayed instant quenching, fast quenching and slow quenching. Instant quenching is defined as particles with YFP-Vpr signal dropping to the background level immediately (≤16 sec) upon adding a low pH buffer. Delayed instant quenching is defined as YFP-Vpr signal quickly dropping after a lag of more than 30 sec following low pH exposure.

Slow YFP quenching, likely resulting from the baseline proton permeability of the viral membrane, was defined based upon gradual quenching of YFP in coverslip-adhered pseudoviruses that do not have viral glycoprotein responsive to low pH. The average slow YFP quenching curve was used to define slow vs fast individual quenching events based upon the intensity traces, respectively, falling within or outside the standard deviation range of the mean YFP loss curve.

### Soluble LAMP1 expression and purification

Soluble LAMP1 (sLAMP1) expression and purification was described previously [32]. Briefly, Expi293F cells were transfected with pHLsec-LAMP1 fragment, a kind gift of Juha T. Huiskonen (University of Oxford) and incubated for three days at 37 °C in the presence of 1 μg/ml of kifunensine (R&D Systems, MN, USA). Supernatant was collected and combined with half a volume of binding buffer (25 mM HEPES, 150 mM NaCl, pH 7.2). His-tagged soluble LAMP1 fragment was purified by Ni-NTA affinity chromatography. The elution was desalted and concentrated to 5 mg/ml.

### Virus-cell fusion assay

A549 cells were seeded in phenol red-free DMEM with 10% FBS the day before infection. BlaM-Vpr containing HIV-1 particles (50 pg p24) pseudotyped with wild-type or GPc were bound to cells by centrifugation at 1550xg for 30 min, 4 °C. Cells were washed with cold phenol red-free DMEM with 10% FBS and 20 mM HEPES (GE Healthcare Life Sciences) to remove unbound viruses and incubated at 37 °C to initiate virus-cell fusion. Cells were then placed on ice, loaded with the CCF4-AM BlaM substrate (Life Technologies), and incubated at 11 °C overnight to allow substrate cleavage. The cytoplasmic BlaM activity (ratio of blue to green fluorescence) was measured using a Synergy H1 plate reader (Agilent Technologies, Santa Clara, CA, USA).

## Acknowledgements

The authors wish to thank Drs. Cohen-Dwashi and Ron Diskin for certain LASV GPc mutants, Dr. Kosuke Miyauchi for cloning the Gag-EcpH construct, Zalgen Labs for a kind gift of 12.1F anti-Lassa GPc antibody. We are indebted to the Melikian lab members, Mariana Marin, Gokul Raghunath, Manish Sharma, Monica Cortez, and David Prikryl for critical reading of the manuscript and helpful suggestions.

## Supplemental Figure Legends

**Figure S1. IAVpp fusion with A549 cell.** (A) Illustration of mCherry-CL-YFP-Vpr labeled single IAVpp fusion. IAVpp is internalized and trafficked to acidic endosomes where it fuses with the endosomal membrane without prior membrane permeabilization (YFP quenching). IAVpp-endosome fusion results in mCherry release into the cytoplasm. (B) A single IAVpp fusion event in A549 cell. Time-lapse images (top) and fluorescence traces (bottom) show virus fusion (mCherry loss) at 1.6 min (see Movie S2).

**Figure S2. Efficiency of “leaky” single LASVpp and MACVpp fusion in different cell lines.** HIV-1 based pseudoviruses pseudotyped with LASV or MACV GPc and labeled with mCherry- CL-YFP-Vpr were attached to cells by spinoculation in the cold, and their entry/fusion was triggered by shifting to 37 °C. Percent of particles releasing mCherry after the YFP-Vpr signal quenching is plotted. Data are means ± SD of 3 independent experiments. Results were analyzed by Student’s t-test. **, p<0.01; NS, not significant.

**Figure S3. Efficiency and kinetics of arenavirus VLP fusion.** (A) Fusion efficiency of single Candid-1, LASV, JUNV and MACV VLPs. Data shown are means ± SD of 3 independent experiments. Results were analyzed by Student’s t-test. *, p<0.05; NS, not significant. (B) Kinetics of fusion of Candid-1, LASV, JUNV and MACV VLPs.

**Figure S4. Nearly all LASVpp limit diffusion of protons through the virion membrane.** (A) Images of coverslip-adhered LASVpp in PBS (left) and 16 seconds after applying membrane-impermeable citrate pH 5.0 buffer (right). (B) Normalized YFP intensity of the only two particles that undergo instant YFP quenching and two representative particles that limit proton diffusion across their membranes. The point of adding a low pH buffer is marked with a red arrow. Tracked particles in (B) are marked by color-matched arrows in (A). Instant YPF quenching events constitute 3.9% of all particles. As shown in Fig. 4, the YFP signal from the rest of single pseudovirions gradually decays over the course of several minutes (not noticeable on a short time scale shown in panel B), due to a baseline proton diffusion or due to increased membrane permeability associated with GPc refolding.

**Figure S5. LASV-GPc efficiently incorporates into HIV-1 particles irrespective of virus labeling approaches.** (A) Images of LASVpp labeled with mCherry-YFP-Vpr or pHuji/Gag- EcpH. Pseudoviruses were bound to poly-L-lysine coated coverslips, fixed, and incubated with anti-LASV GPc human antibody, followed by staining with anti-human AF647-conjugated antibody. Viral mCherry or EcpH markers were used to identify viral particles and visualize the associated GPc signal. (B) Quantification of co-localization of LASV GPc immunofluorescence with viral particles identified by mCherry or EcpH fluorescence, as indicated. (C) Distributions of the GPc immunofluorescence intensities for each virus.

**Figure S6. Time lapse images at indicated time points after adding low pH buffer to viruses on coverslips and on cells.** YFP fluorescence quenching at low pH is readily detectable, while the reference mCherry signal is largely unchanged. The apparent loss of mCherry puncta on A549 cells (lower panel) is due to cell shrinkage at low pH which results in a fraction of particles moving out of focus.

**Figure S7. Examples of four types of YFP-Vpr quenching of single LASVpp on coverslip (A-D) and on DF-1 cells (E-H).** The time points when low pH buffer was added are marked with red arrows. See Fig. 4.

**Figure S8. Estimation of viral membrane permeability to protons.** Representative pseudoviruses exhibiting fast and slow YFP quenching were used to estimate their membrane permeability to protons using the equation P=k*V/S, where k is the permeability coefficient determined by exponential fit of normalized YFP fluorescence decay (assuming that fluorescence changes reflect changes in intraviral proton concentration), V is the volume and S is the surface area of the virus. The P values shown are for slow and fast quenching events assuming the particle radius is 60 nm and that the enclosed inner volume is “empty”, which likely results in overestimation of the permeability value. Nonetheless, the obtained P values are much lower than those reported in the literature for liposomes. It is thus possible that the viral interior proteins buffer the inner pH by binding the incoming protons and slowing down YFP quenching.

**Figure S9. Viral glycoproteins are efficiently incorporate into HIV-1-based pseudoviruses.** (A) Images of the pseudoviruses labeled with mCherry-CL-YFP-Vpr. Pseudoviruses were bound to poly-L-lysine coated coverslips, fixed and incubated with anti- LASV-GPc, anti-HIV Env or anti-IAV HA antibodies, as indicated in the figure. Viruses then were stained with respective AF647-conjugated secondary antibodies. Viral particles were identified based on the mCherry signal. Bald particles lacking envelope glycoproteins and GPc-containing viruses produced in the presence of saquinavir viruses were used as controls. (B) Quantification of co-localization of viral glycoproteins detected by immunofluorescence signal and viral particles labeled with mCherry. (C) Distribution of GPc immunofluorescence intensity for different pseudoviruses.

**Figure S10. LASV receptors, α-dystroglycan and LAMP1, do not promote LASVpp membrane permeabilization.** LASVpp were bound to the cell surface or to poly-L-lysine coated coverslips at 4 °C. GPc conformational changes are triggered by applying membrane impermeable pH 5.0 citrate buffer. (A) For soluble LAMP1 (sLAMP1) treatment, 200 µg/ml of sLAMP1 was added to viruses in a pH 5.0 citrate buffer. For LAMP1 antibody (LAMP1ab) treatment, cells were incubated with the growth medium containing 100x diluted LAMP1ab for 1 hour before imaging; LAMP1ab was also present in the pH 5.0 citrate buffer. Two hundred µg/ml of BSA was in citrate pH 5.0 buffer used as a control. (B) For recombinant human α- dystroglycan (rhDG) treatment, LASVpp were incubated with 50 µg/ml of rhDG or BSA (control) at 37 °C for 20 min, the pH 5.0 citrate buffer was supplemented with 50 µg/ml of BSA or rhDG. The rate constants k are in 1/sec.

**Figure S11. Mutant LASV-GPc efficiently incorporate into LASVpp.** (A) Images of the pseudoviruses labeled with mCherry-CL-YFP-Vpr. Pseudoviruses were bound to poly-L-lysine coated coverslips, fixed, and incubated with anti-LASV GPc human antibody, followed by staining with AF647-conjugated anti-human antibody. Viruses were identified based on the mCherry marker. (B) Quantification of the co-localization of GPc immunofluorescence and viral particles identified by mCherry. (C) Distribution of the GPc immunofluorescence intensities for each virus.

## Movie legends

**Movie S1. Single LASVpp fusion with A549 cells manifested in sequential YFP quenching and dequenching.** Single LASVpp exhibits YFP quenching at 12.3 min followed by YFP dequenching/mCherry loss at 25.6 min corresponding to virus interior acidification and fusion, respectively. Movie is related to Fig 1B.

**Movie S2. Single IAVpp fusion with A549 cells manifested in mCherry release.** Single IAVpp exhibits mCherry loss at 1.6 min corresponding to virus fusion. Movie is related to Fig S1B.

**Movie S3. Single Candid-1 VLP fusion with A549 cells manifested in sequential YFP quenching and dequenching.** Single Candid-1-VLP exhibits YFP quenching at 12.1 min followed by YFP dequenching at 33.4 min corresponding to VLP interior acidification and fusion, respectively. White arrows marked the VLP of concern. Movie is related to Fig. 2B.

**Movie S4. Single LASVpp enters acidic compartments and undergoes membrane permeabilization in A549 cells.** Single LASVpp exhibits pHuji quenching at 107.6 min followed by EcpH quenching at 108.2 min corresponding to virus entering acidic endosome and interior acidification, respectively. Movie is related to Fig. 3B.

**Movie S5. Single IAVpp enters acidic compartments without membrane permeabilization in A549 cells.** Single IAVpp exhibits pHuji quenching at 23.9 min corresponding to virus entering acidic endosome. Movie is related to Fig. 3C.

